# Recombinant measles virus equipped with BNiP3, a pro-apoptotic gene, targets β-catenin pathway in triple negative breast cancer cells

**DOI:** 10.64898/2026.04.15.718830

**Authors:** Alok Kumar, Kamaldeep, Gauri Shankar Upadhyay, Maitreyi S. Rajala

## Abstract

Oncolytic virotherapy is an emerging cancer therapy using genetically modified viruses. We previously reported engineering of measles virus with BNiP3, a proapoptotic gene for oncolytic purposes. The recombinant virus had shown promising results in breast cancer cells with a bias towards TNBC, an invasive and an aggressive subtype. Here, we investigated the mechanistic insights of anti-tumor effects induced by the recombinant virus. Initially, TNBC and non-TNBC tumor cell lines were compared bioinformatically using the available gene expression data through protein-protein interaction network using different topological properties. Four hub genes involved in tumor development and progression were identified to be the top genes in both the data sets. Of which, CTNNB1 gene encoding β-catenin was found to be the significant one; as β-catenin pathway is known to be a driver of tumor cell invasion and migration, the impact of the virus on this pathway was investigated in breast tumor cells. The results had demonstrated a notable decrease in β-catenin expression and its downstream targets, cyclin D1, MMP7 reducing the migration potential of TNBC cells following virus infection. These findings suggest that the recombinant measles virus could be one of the effective treatment modalities to target invasive TNBC tumors. *In vivo* validation of engineered virus is ongoing to explore the therapeutic application of this virus.

**Graphical Abstract:** 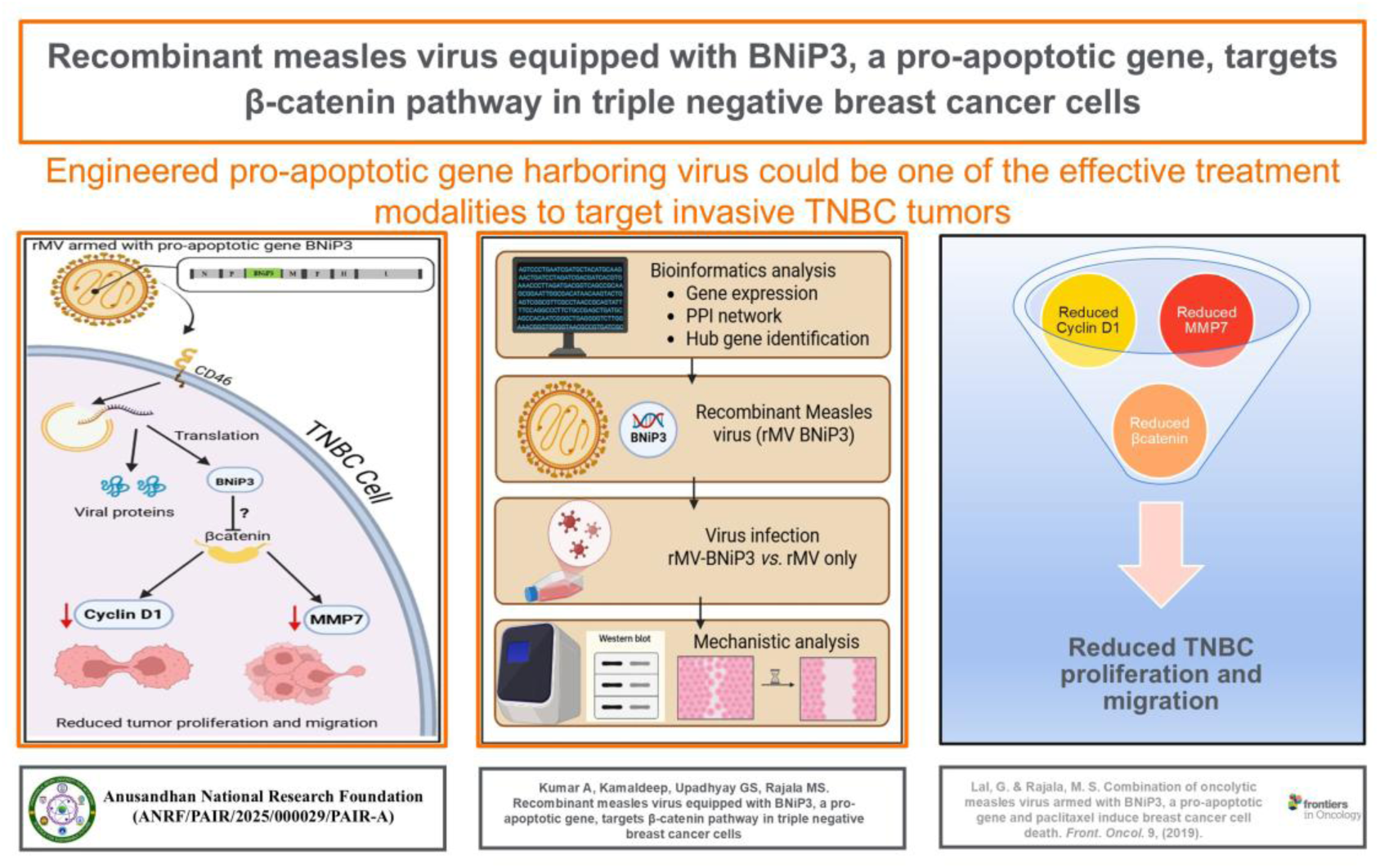

**Highlights:**

- Use of recombinant measles virus with a pro-apoptotic gene, BNiP3 to target breast cancer cells
- Identification of top regulatory genes in breast cancer development and progression
- Reduction of β-catenin expression encoded by CTNNB1 gene in TNBC cells following virus infection
- Downregulation of β-catenin downstream targets in TNBC cells with virus infection
- Inhibition of migratory potential of TNBC cells following infection

## Introduction

Triple-negative breast cancer (TNBC) is a diverse subset of breast cancer that accounts for about 15–20% of all breast cancer cases ^1^. It is predominantly found in women under the age of 40 years and in the African-American ethnicity ^2^. TNBC is an aggressive subtype by nature and commonly observed as infiltrating ductal carcinoma, but it can also be medullary-like cancers, metaplastic cancers, and rare forms such as adenoid cystic carcinoma ^3^. Notably, almost 80% of TNBC cases are linked to BRCA1 mutations ^4^ whereas only a few cases may be associated with BRCA2 mutations ^5^.

Characteristics of TNBC include a high level of cell invasion and metastasis to other organs such as the brain, lungs, and liver, resulting in poor outcome and reducing the chances of patient survival ^6^; ^7^. The lack of estrogenic receptor (ER) and progesterone receptor (PR) expression, combined with the absence of HER2 expression and the significant heterogeneity of TNBC has posed challenges in developing effective treatment options for this breast cancer subtype. TNBC treatment typically includes surgery and chemotherapy, which are often unsuccessful, leading to recurrence or metastasis. Besides, numerous other strategic therapeutic approaches have been formulated to address TNBC subtypes, but limitations still remain. Since there is no single treatment option that is effective against all subtypes of TNBC, there is a need for a newer, efficient therapeutic approaches involving multimodal application of therapies in synergistic combination.

Oncolytic virotherapy (OV) is an innovative treatment approach that involves the genetic engineering of viruses to specifically target cancer cells while sparing nearby non-cancerous cells. Measles virus (MV) is one potential virus to trigger tumor regression, which can be manipulated for better onco-therapeutic purposes. This idea led to the approval of the vaccine strain for cancer therapy back in 1960 ^8^. Our lab earlier engineered measles virus by arming with BNiP3, a proapoptotic gene of human origin for oncolytic purpose using the vaccine strain of measles virus full-length genome backbone (plasmid vector pMV-NSE-FlagP-M502-3p was a gift from Branka Horvat). We had reported that the recombinant measles virus harboring BNiP3 gene (rMV-BNiP3) induced anti-proliferative effects in breast cancer cells with better selectivity towards MDA-MB-231 cells, a representative cell line of TNBC. MDA-MB-231 cells infected with the rMV-BNiP3 virus had shown significant apoptosis compared to MCF-7, a non-TNBC cell line ^9^. However, the molecular mechanism that contributes to the cell death following oncolytic measles virus infection is an important aspect to be elucidated.

The current study was designed to understand the mechanistic insights of the engineered measles virus, rMV-BNiP3-induced cytotoxicity/cell death in TNBC cells. Prior to *in vitro* experiments, the key drivers in TNBC development and progression were identified using bioinformatics tools with the available gene expression data of both the TNBC and non-TNBC cell lines. This analysis was done to see if rMV-BNiP3 would have an impact on any of the identified major drivers of TNBC. The CTNNB1, a gene encoding β-catenin was found to be one of the key regulators of TNBC development and progression. The gene expression profile was validated at the transcript level in MDA-MB-231 *vs*. MCF-7, representative cell lines of TNBC and non-TNBC respectively. As β-catenin is known to be upregulated in various invasive tumors, including invasive TNBC, the impact of oncolytic rMV-BNiP3 infection on β-catenin pathway and its downstream targets was evaluated. Most of the screening methods have been performed in 2D monolayer cell cultures while 3D spheroid cell cultures were used to check the migration potential of TNBC *vs*. non-TNBC cells following infection. Taking the bioinformatics and the experimental conclusions together, our findings indicate that the recombinant measles virus armed with pro-apoptotic gene, BNiP3 (rMV-BNiP3) has a potential to target β-catenin pathway and may suppress the proliferation and invasion of aggressive breast tumor subtype, TNBC.

## Materials and Methods

### Cell cultures and maintenance

MDA-MB-231(invasive breast carcinoma) and MCF-7 (non-invasive breast carcinoma) cell lines were purchased from the National Centre for Cell Science (NCCS), Pune. They were maintained in Dulbecco’s modified eagle’s medium (DMEM) containing 10% FBS and antibiotics at 37 °C in 5% CO2 incubator.

### Recombinant oncolytic measles virus

A genetically modified measles virus generated earlier in our lab for the oncolytic purpose with BNiP3, a pro-apoptotic gene of human origin (rMV-BNiP3), was used in this study ^9^. Recombinant measles virus without BNiP3 gene (rMV) was used for comparison in a few experiments.

### Bioinformatics studies

For analysis, the datasets of TNBC *vs*. non-TNBC cell lines; more specifically, MDA-MB-231 and MCF-7 breast cancer cell lines were chosen, and the gene information was retrieved from the database (gene expression data) maintained by Rajiv Gandhi Centre for Biotechnology, India ^10^. These two breast cancer cell lines were selected for validation and comparison in the current study. In order to generate the protein-protein interaction (PPI) network, the interconnected genes from cell line level analysis were extracted from the STRING database ^11^, followed by visualization using Cytoscape version 3.8.2 ^12^ Cytohubba ^13^ and the Network Analyzer plugin of Cytoscape were utilized to calculate topological properties of the PPI network. For the identification of the hub genes through PPI network, multiple topological properties were considered. Initially, the topological properties such as highest degree, bottleneck, and betweenness centrality were used to determine hub genes that are differentially expressed between MDA-MB-231 and MCF-7 cells. Further, the key hub genes were identified using ten algorithms: betweenness centrality, bottleneck, closeness centrality, degree centrality, eccentricity, stress, radiality, maximum neighbourhood component (MNC), maximum clique centrality (MCC), and edge percolated component (EPC).

### *In vitro* studies Virus infections

The key regulatory genes identified by bioinformatics tools were validated in breast cancer cell lines infected with purified virus. The recombinant measles viruses (rMV and rMV-BNIP3) were purified using the iodixanol density gradient method ^14^. MDA-MB-231 and MCF-7 cells were grown in 35 mm dishes to 90% confluency. Cells were infected with purified virus (rMV or rMV-BNiP3) and mock infected with DMEM. Infections were done in two sets and incubated in a 37°C incubator with 5% CO₂ for 24 to 48 hours till the visualization of cytopathic effect (CPE). At 48 hours post-infection, cells were harvested; total RNA was isolated from one set of cells and used for quantitative real-time PCR analysis. Another set of infected cells was also harvested, from which total protein was recovered for SDS-PAGE and western blot analysis.

### Quantitative real time PCR

The expression of top hub genes identified to be the potential key regulators of breast tumorigenesis by bioinformatics tools was validated in invasive/TNBC and non-invasive/non-TNBC (MDA-MB-231 and MCF-7) cell lines at the transcript level by quantitative real-time PCR using specific primers. Total RNA was isolated from cells by the Trizol method (Thermo Fisher Scientific), followed by cDNA synthesis using Revert Aid Reverse Transcriptase Enzyme (Thermo Fisher Scientific). A fraction of cDNA was subjected to quantitative real-time PCR using specific primers to the CTNNB1 (FP: 5’ CACAAGCAGAGTGCTGAAGGTG 3’, RP:5’GATTCCTGAGAGTCCAAAGACAG 3’); c-JUN (FP: 5’CCTTGAAAGCTCAG AA CTCGGAG 3’, RP: 5’ TGCTGCGTTAGCATGAGTTGGC 3’); ERBB2 (FP: 5’CCTTGA AAG CTCAGAACTCGGAG 3‘, RP: 5’TGCTGCGTTAGCATGAGTTGGC 3’); CDH1 (FP: 5’GCC TCCTGAAAAAGAGAGT GGAAG 3’, RP: 5’TGGCAGTG TCTCTCCA AATCCG 3’); and GAPDH (FP: 5‘TGCACCACCAACTGCTTAGC 3’, RP: 5‘GGCATGGACTGTGGTCATG AG 3‘) by the SYBR green method. Primer sequences were taken from published papers from Origene website (www.origene.com). Each sample was run in technical triplicates and the mean Ct value was taken for each set of reactions. The comparison of two samples was carried out by the determination of the ’n’ fold difference in mRNA abundance through relative quantification using the 2^-ΔΔCt^ method.

### SDS-PAGE and western blot analysis

MDA-MB-231 or MCF-7 cells were infected with rMV (recombinant measles virus) or rMV-BNiP3 (recombinant measles virus with BNiP3 gene). At 24 hours infection, the cells were harvested, and the total protein was recovered by adding 1 mL of RIPA lysis buffer containing 20 mM Tris-HCl, pH 7.5, 150 mM NaCl, 5 mM EDTA, 1% NP-40, 0.5% sodium deoxycholate, 1 mM PMSF, 1 mM sodium orthovanadate, and a protease inhibitor cocktail. The protein concentration was determined by Bradford reagent. Equal amount of protein from each sample was subjected to SDS-PAGE followed by western blot analysis using specific antibodies against BNiP3, CD46, β-catenin, Cyclin D1, MMP7, c-JUN, phospho-and total AKT, phospho-Erk2, and β-actin (Cell Signaling Technology Inc.). Densitometric analysis of the target protein bands was performed using ImageJ software, and the band intensity was normalized against its appropriate loading control.

### Wound healing assay

The migration potential of breast cancer cells in the presence or absence of recombinant virus was checked by wound healing assay. MDA-MB-231 and MCF-7 cells were cultured in 12-well plates. When the cells reached to 80-90% confluency, a scratch was made with a 200 µL pipette tip. Cells were then infected with rMV or rMV-BNiP3 and incubated for 48 hours. Mock infected cells were used as a control. At 48 hours infection, the migration potential of infected and mock-infected cells was observed under a phase contrast microscope at 10X magnification, and the images were captured.

### Development of 3D spheroid cultures

Spheroids were created using a hybrid approach that combined both hanging drop and agarose base methods. The additives, methocel and collagen were included to enhance the circularity and compactness, resulting in improved spheroid formation. MDA-MB-231 and MCF-7 cells were grown in a 25 cm² flask until they reached approximately 70-80% confluence. Then, the cells were trypsinized and centrifuged, and the pellet was resuspended in an incomplete medium. The concentration was adjusted to 1-5 x 10^5 cells/ml and placed onto the lid of a plate and inverted upside down. Dilutions were made to adjust 100-500 cells. The minimum volume taken for seeding was 10 µl. Sterile petri dishes were coated with 1-1.5% of agarose, and the agarose was allowed to solidify at room temperature. Agarose was overlaid with 200 µl of complete medium, and the spheroid formed by the hanging drop method was carefully transferred onto the agarose plate. Plates were then incubated at 37°C with 5% CO₂ till the spheroids grew to considerable volume. Then, the MDA-MB-231 and MCF-7 cell spheroids were infected with rMV or rMV-BNiP3. Spheroids are checked every day, and the images were captured every 2 days.

## Results

### Identification of key regulators in breast tumor development and progression

Analysis of TNBC and non-TNBC cell line data using bioinformatics tools on the basis of topological properties had shown prominent genes involved in TNBC development and progression. Based on the first three algorithms— highest degree, bottleneck and betweenness centrality—CTNNB1, CDH1, ERBB1, and c-JUN genes were found to be the top key regulators in TNBC development (Fig 1a). Further analysis to identify the key regulators in specific cell lines, MDA-MB-231 (TNBC) *vs*. MCF-7 (non-TNBC) using all ten algorithms also had shown CTNNB1 and CDH1 as top genes in concurrence with the data obtained using three algorithms (Suppl Fig 1). The genes that were identified to be top key players were validated by quantitative real-time PCR. The CTNNB1 gene encoding β-catenin was found to be overexpressed, and the downregulation of CDH1 was noted in MDA-MB-231 cells in comparison with MCF-7 cells. While the other two genes, EGFR1 shown not much difference between two cell lines used while JUN, were also found to be highly expressed in MDA-MB-231 cells (Fig 1b). The CTNNB1 gene was found to be the top hub gene as shown by using either three or ten algorithms, indicating its encoding protein β-catenin is a key regulator of TNBC development and progression as reported in the literature.

**Fig 1.**
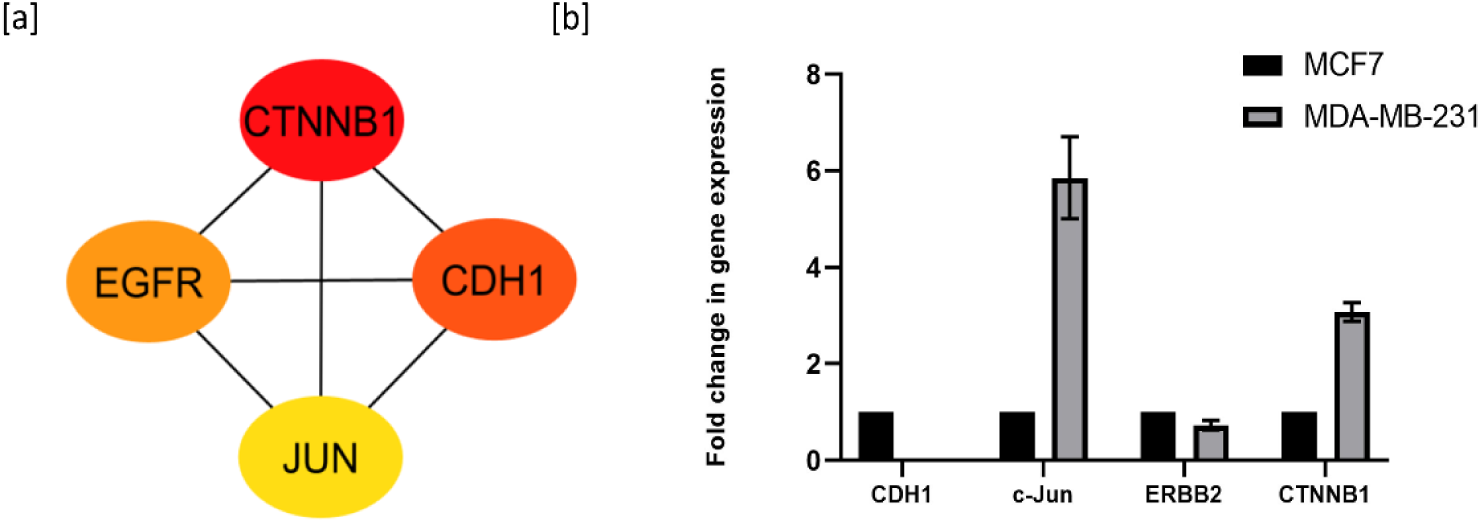
Identification of hub genes, key regulators of TNBC development and progression. The datasets of TNBC and non-TNBC (MDA-MB-231 and MCF-7) cell lines were chosen for the identification of the key regulators in tumor development and progression. Hub genes were identified through PPI network using three topological properties, highest degree, bottleneck and betweenness centrality. [a] Sub network construction of differentially expressed top hub genes between MDA-MB-231 *vs*. MCF-7. Red color indicates upregulated genes [b] *In vitro* validation of identified top key regulators in MDA-MB231 *vs.* MCF-7 cells by quantitative real time PCR.

### Confirmation of purified recombinant measles virus infection in TNBC cells

The replication efficiency of purified rMV and rMV-BNiP3 was confirmed in both the cell lines. The TNBC cell line had shown better CPE compared to the non-TNBC cell line. Viral ‘N’ gene encoding nucleoprotein expression in rMV- and rMV-BNiP3-infected cells confirmed the infection (Fig 2a). The foreign gene, BNiP3 expression following rMV-BNiP3 infection in TNBC cells at the transcript (Fig 2b) and protein level (Fig 2c) was confirmed. The expression of CD46, the receptor being used by the measles virus vaccine strain (Edmonston strain, the virus that was used for the development of rMV) was compared between MDA-MB-231 and MCF-7 cells in order to determine the cause of the bias of the engineered virus towards TNBC cells. Slightly higher expression of CD46, but not significant difference was noted in MDA-MB-231 cells (Fig 2d). Densitometric analysis of the blot shown in Fig 2e.

**Fig 2.**
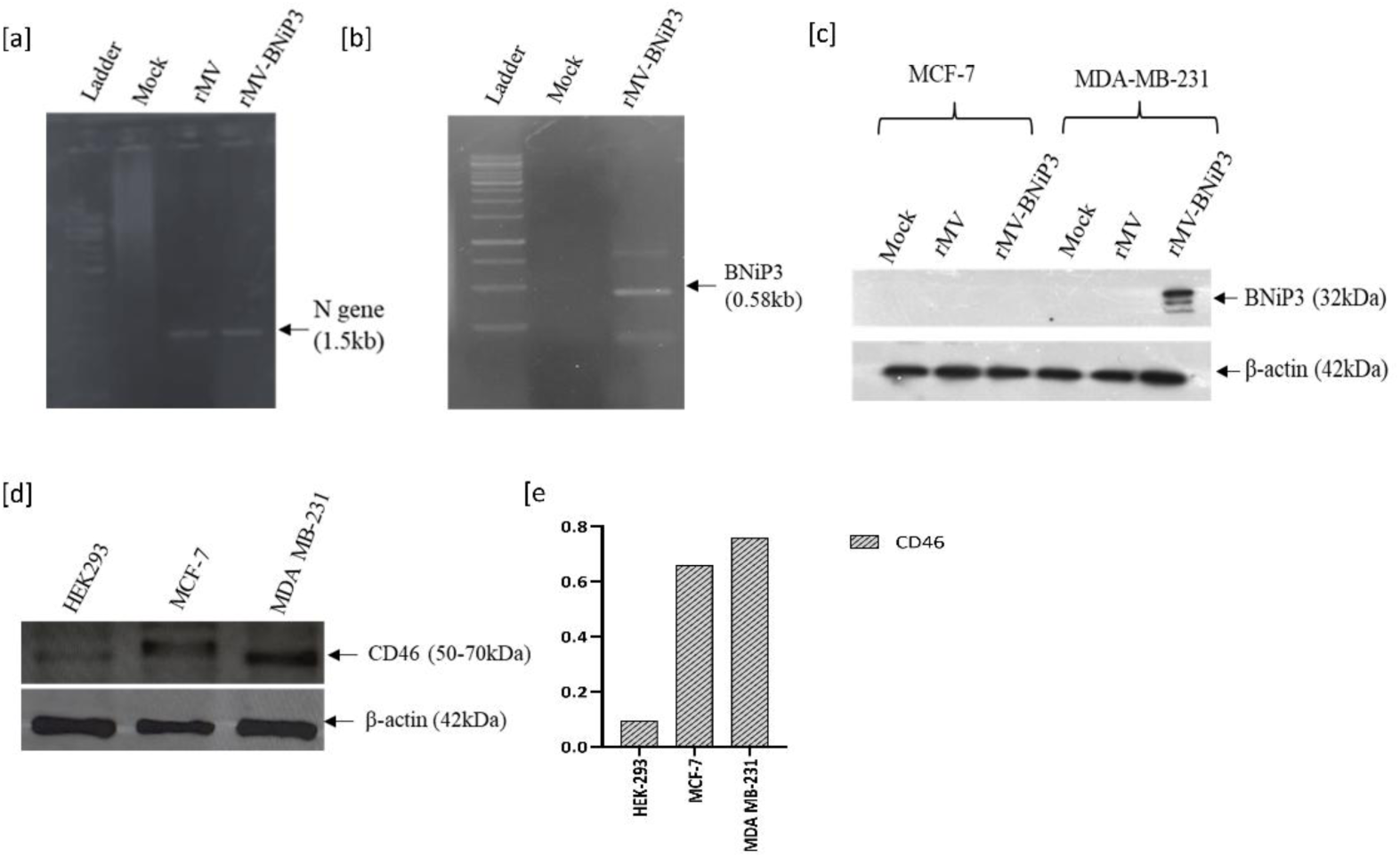
Recombinant measles virus without BNiP3 (rMV) and with BNiP3 (rMV-BNiP3) infection in breast tumor cells. MDA-MB-231 cells were infected with purified rMV or rMV-BNiP3 and the expression of viral and inserted gene was confirmed by real time PCR and western blot analysis. [a] Viral gene encoding nucleoprotein expression in infected cells. Expression of BNiP3 at [b] transcript [c]protein level [d] Expression of CD-46, the receptor being used by measles virus vaccine strain in both the cell lines. HEK293 was used for comparison [e] densitometric analysis of CD46 blot.

### Reduction of β-catenin expression

In TNBC, β-catenin-dependent Wnt Signaling is necessary for cell migration, invasiveness and is associated with reduced overall survival in breast cancer patients ^15^. The aberrant activation of this pathway enhances stem cell characteristics in TNBC cells. Thus, the impact of oncolytic measles virus, rMV or rMV-BNiP3 infection on β-catenin pathway was evaluated. The expression of β-catenin in MDA-MB-231 cells was almost reduced to no expression following infection with the rMV-BNiP3 as compared to cells infected with rMV alone (Fig 3a). Densitometric analysis of the western blot was also shown (Fig 3b). No such substantial reduction of β-catenin expression was observed in MCF-7 cells following infection.

**Fig 3.**
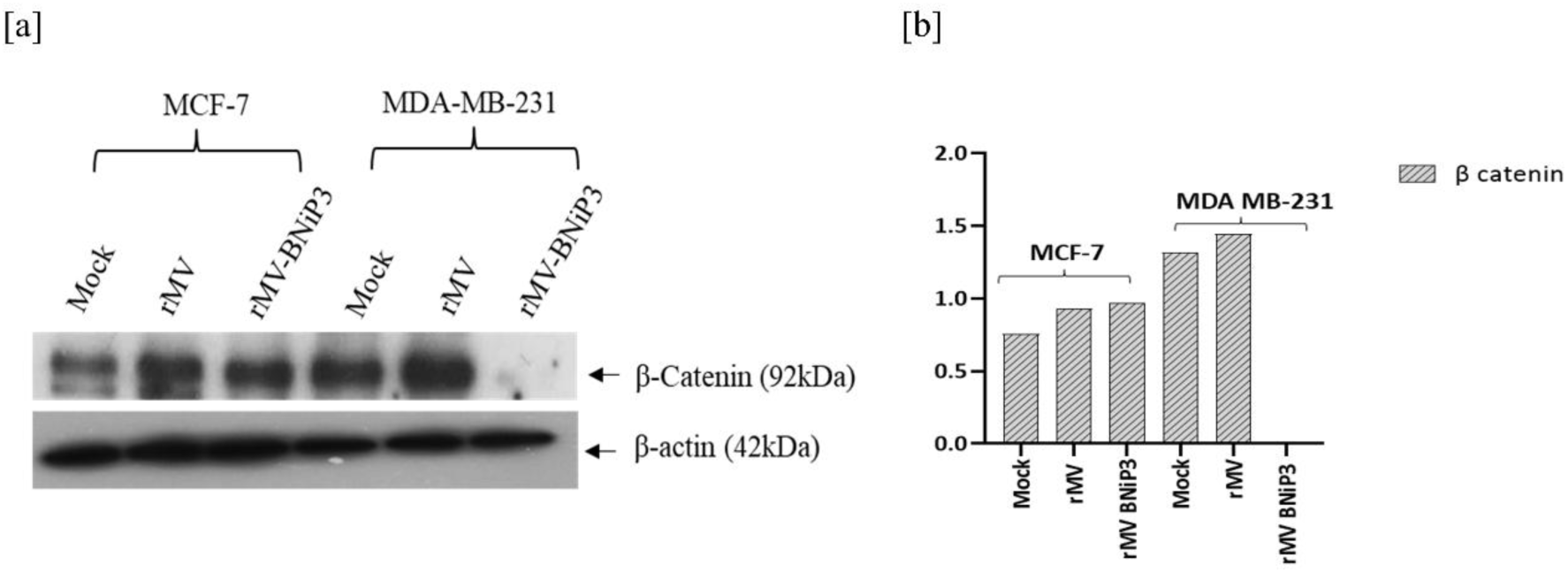
Regulation of β-catenin expression. MCF-7 and MDA-MB-231 cells were infected with rMV or rMV-BNiP3 in duplicates. At 48 hours post infection, total protein was recovered, quantified and equal amount of protein from each sample was subjected to SDS-PAGE followed by western blot analysis using anti β-catenin antibody. Mock infected cells were used as a control. [a] Expression of β-catenin [b] Densitometric analysis

### Inhibition of downstream targets of β-catenin pathway

The expression of β-catenin downstream targets, Cyclin D1, MMP7, and c-JUN, that are involved in tumor progression and invasion was evaluated in MCF-7 *vs*. MDA-MB-231 cells infected with rMV or rMV-BNiP3. Cyclin D1, a major regulator of the cell cycle in cancer progression, was noted to be reduced prominently following virus treatment in the TNBC cells over non-TNBC cells. A similar effect was observed on the expression of MMP7 (matrix metalloproteinase-7), a downstream target gene of β-catenin/TCF-4. MMP7 is overexpressed in various cancers including TNBC and is known to be involved in extracellular matrix degradation and tumor metastasis. Following rMV-BNiP3 infection, the MMP7 level was reduced in both the cell lines, but the reduction was more prominent in MDA-MB-231 cells (Fig 4a to 4c). Another downstream target of β-catenin, c-JUN, a regulator of the functional β-catenin–TCF complex, was also reduced following infection (Fig 4d and 4e). Corresponding graphs show the densitometric analysis of each blot normalized with controls. These experiments clearly show that the recombinant measles virus, rMV-BNiP3 downregulates β-catenin and its downstream targets indicating the therapeutic effect on TNBC cells. Interestingly, the anti-tumor effect of the rMV-BNiP3 virus was found to be noteworthy compared to rMV. It appears that BNiP3 is possibly a contributing factor; further research is needed to understand mechanism that is driving these effects.

**Fig 4.**
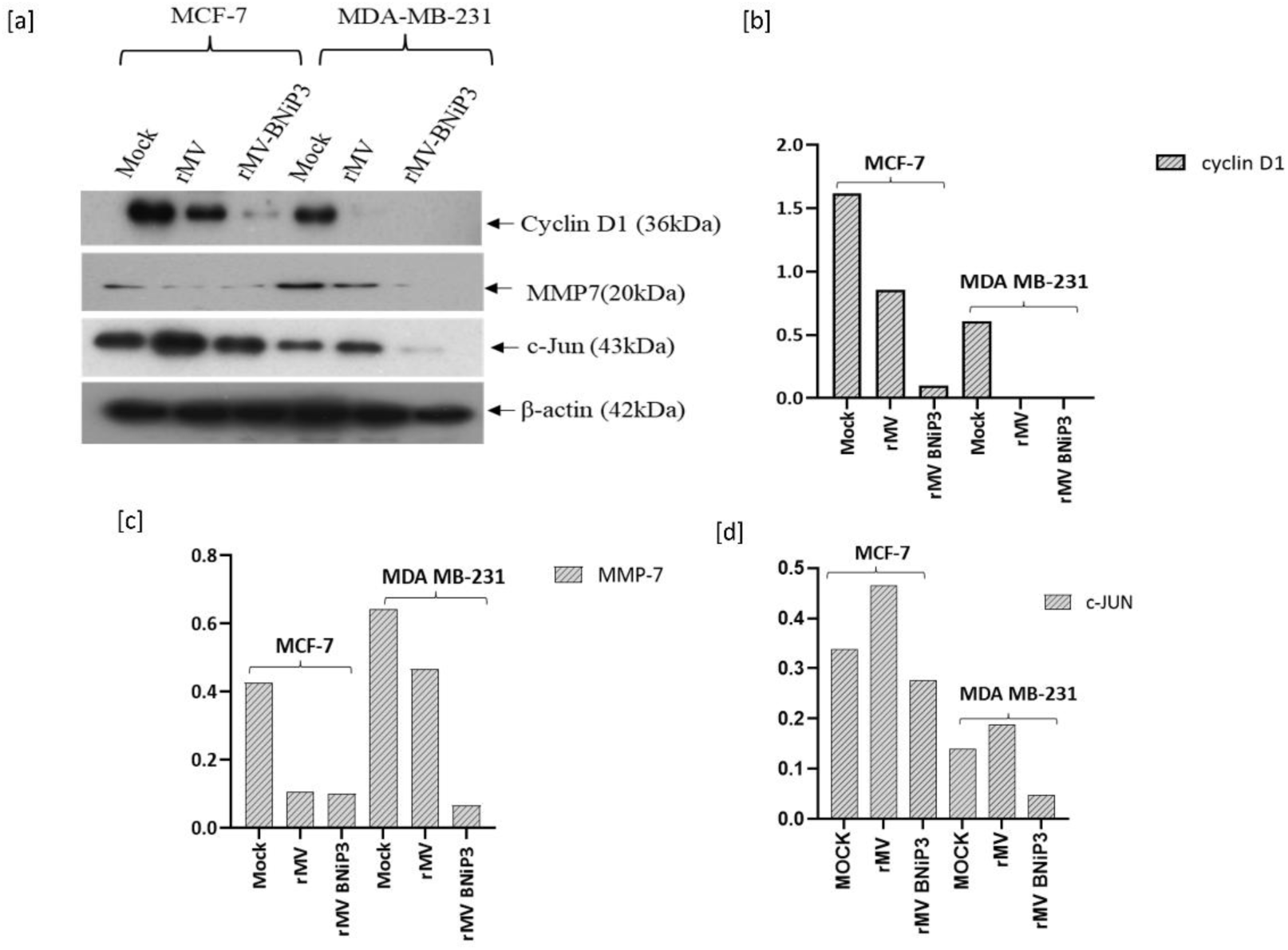
Downregulation of β-catenin downstream targets. MCF-7 and MDA-MB-231 cells were infected with rMV or rMV-BNiP3, and the total protein recovered as described in the previous experiment. At 48 hours post infection, equal amount of protein from each sample was subjected to SDS-PAGE followed by western blot analysis using anti cyclin D1, MMP7 and c-Jun antibodies. Mock infected cells were used as controls. Expression of [a] Cyclin D1, MMP7 and c-Jun. Densitometric analysis of [b] cyclin D1 [c] MMP7 blots [d] c-Jun blots

### Inhibition of proliferative signal pathways

The effect of rMV and rMV-BNiP3 was also evaluated on proliferative pathways, AKT and ERK2, that are largely deregulated in various tumors including breast tumors. PhosphoErk1/2 was observed to be reduced in both MCF-7 and MDA-MB-231 cells infected with rMV-BNIP3 compared to rMV, but the reduction was more apparent in MDA-MB-231 cells (Fig 5a and 5c). Likewise, phosphoAKT level was compared in both the cell lines. Slight reduction of phosphoAKT in MDA-MB-231 cells following rMV-BNiP3 infection was noted although it does not appear to be very significant (Fig. 5b and 5d). Total AKT remained unchanged in both the cell lines and are comparable. β- actin was used to normalize Phospho-Erk1/2. Corresponding densitometric values were shown.

**Fig 5.**
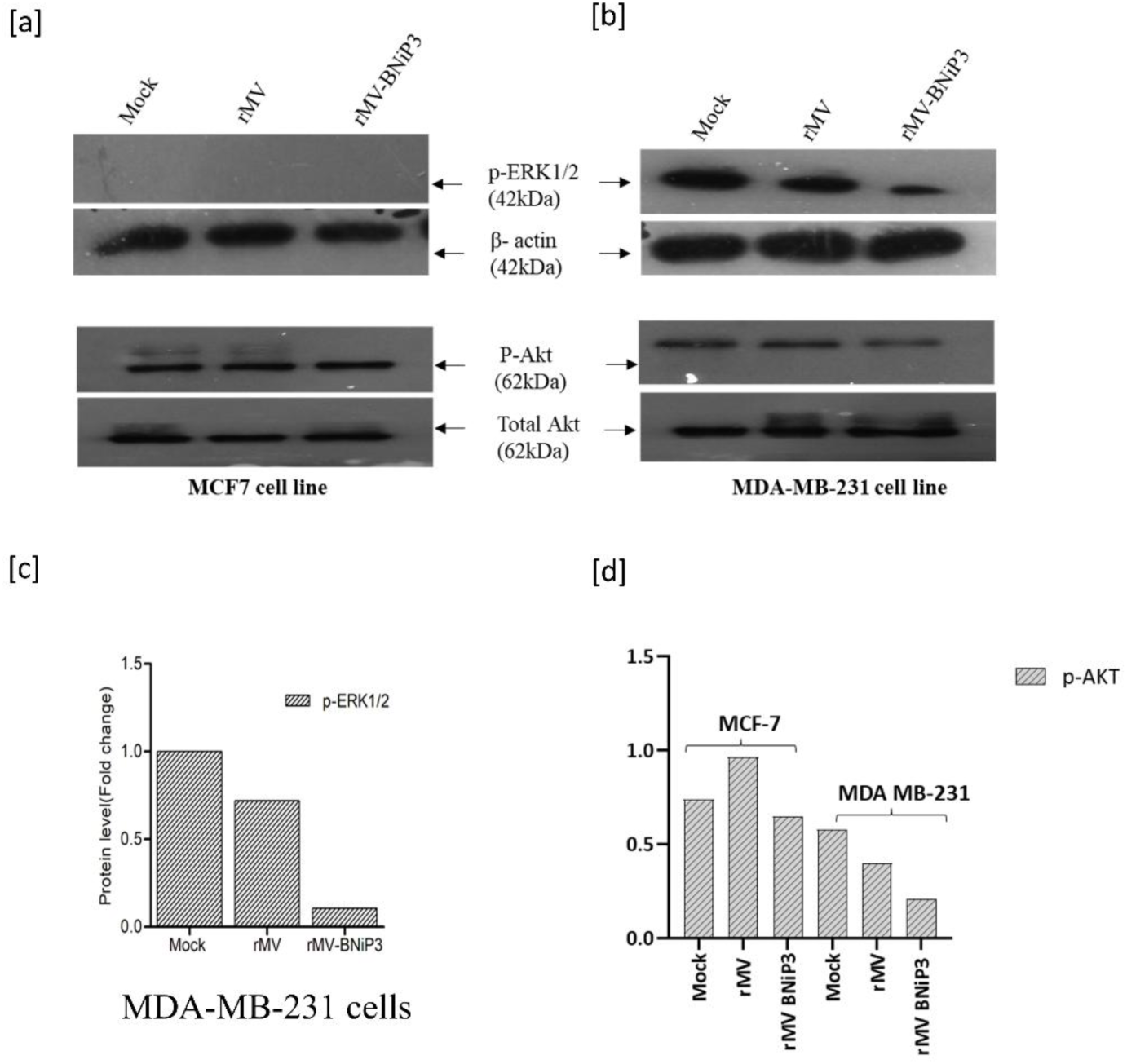
Downregulation of proliferative pathways. Infection and protein recovery was done as described before in MCF-7 and MDA-MB-231 cell. At 48 hours infection, equal amount of protein from each sample was subjected to SDS-PAGE followed by western blot analysis using anti-phospho-Erk1/2, phospho-Akt, and total Akt. Phospho-Erk1/2 and Phospho-Akt level of [a] MCF-7 [b] MDA-MB-231 cells [c] densitometric analysis of ERk1/2 blot [d] densitometric analysis of Akt blot

### Inhibition of tumor cell invasion and migration

It has been widely reported that TNBC tumors are highly aggressive in nature, the growth and migration abilities of MDA-MB-231 and MCF-7 cell lines have been studied by a scratch/wound healing assay. As β-catenin is a key driver of invasion and cell migration, TNBC migration potential was also affected with the reduction of β-catenin and its downstream targets in rMV-BNiP3 infected cells. Visualization under phase-contrast microscope clearly indicate that there is an inhibition in the migration and spreading of TNBC cells compared to mock-infected cells. While no difference in migration was noted in MCF-7 cells before and after infection with the virus (Fig 6). Further, using 3D spheroids, the migration ability of cells infected with virus was evaluated. The rMV-BNiP3 demonstrated anti-proliferative effects, clearly illustrating the differences between the invasive MDA-MB-231 and the non-invasive MCF-7 spheroids (Fig 7). The spheroid size of MDA-MB-231 cells was observed to be reduced following infection, while uninfected cells started forming monolayers by migrating from the spheroid. These finding collectively suggest that the rMV-BNiP3 infection alters the proliferation and migratory capabilities of TNBC cells, potentially impacting their invasive behaviour and overall tumor progression.

**Fig 6.**
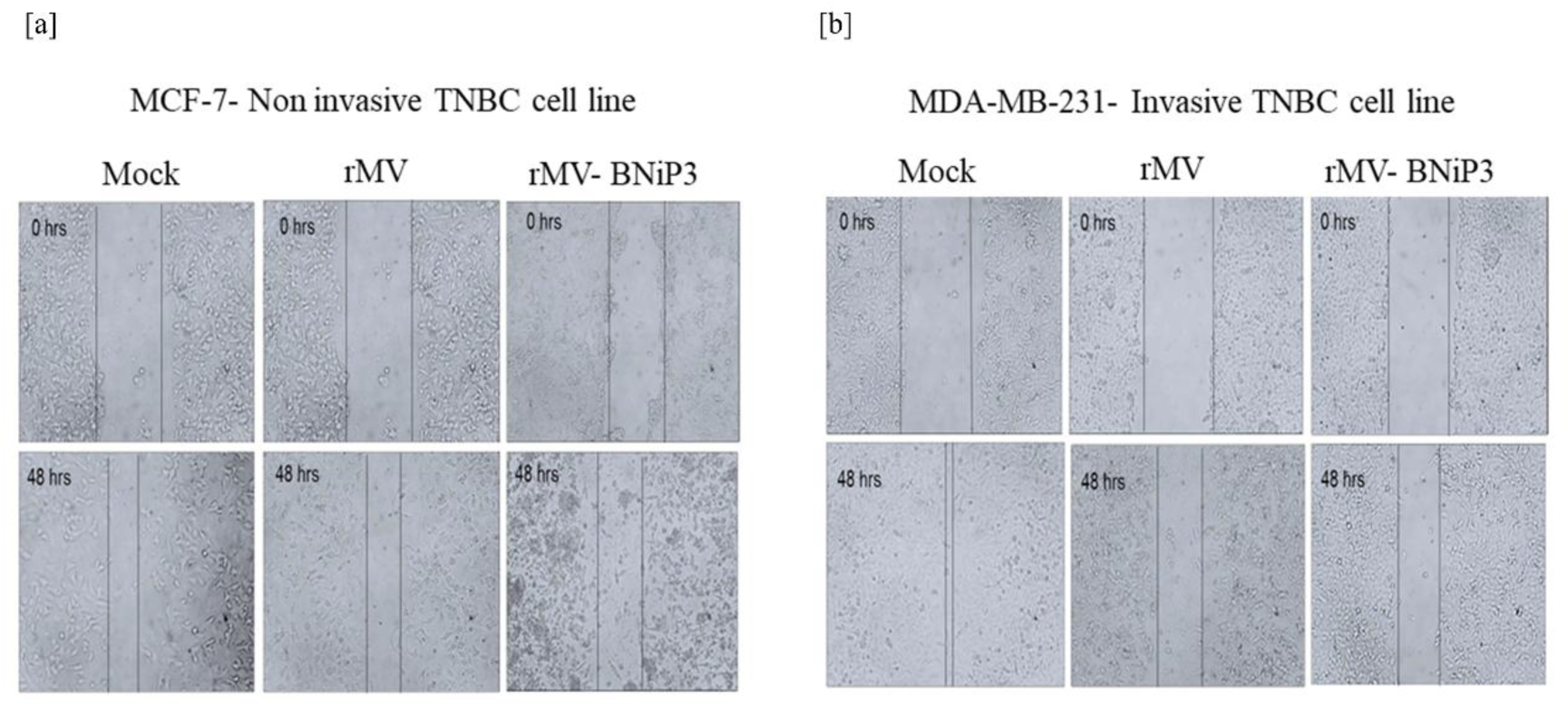
Reduction of migration potential of TNBC cells. MDA-MB-231 and MCF-7 cells were cultured in 12-well plates. After cells reached to confluency, a scratch was made with a 200 µL pipette tip. Cells were then infected with rMV or rMV-BNiP3 and incubated for 48 hours. Mock infected cells were used as a control. [a] MCF-7 cells [b] MDA-MB-231 cells infected with rMV and rMV-BNiP3

**Fig 7.**
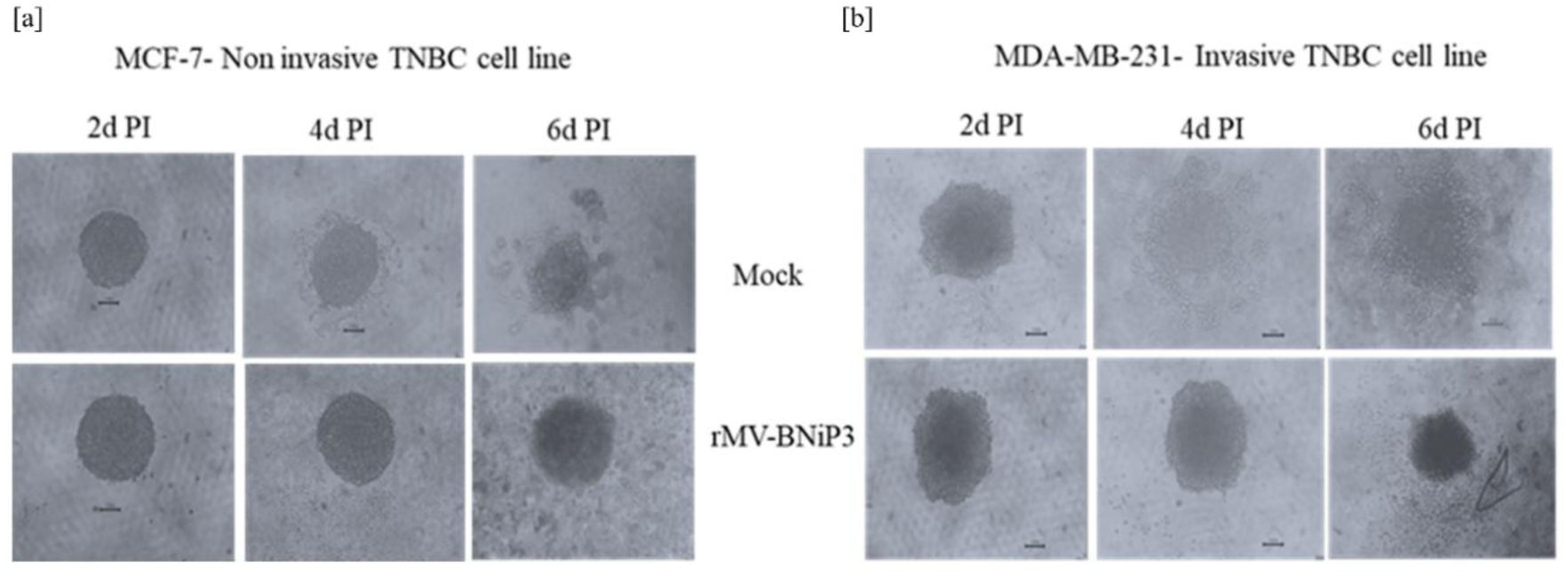
Reduction of spheroid size following rMV-BNiP3 infection. MCF-7 and MDA-MB-231 cell spheroids were created using a hybrid approach that combined both hanging drop and agarose base methods. 3D spheroids were infected with rMV-BNiP3 and are checked every day. Images were captured every 2 days [a] MCF-7 [b] MDA-MB-231 cell spheroids infected with rMV-BNiP3. Mock infected cell spheroids were used as controls

## Discussion

The measles virus possesses unique characteristics that make it a potential therapeutic agent and is currently under study as a novel anticancer agent ^16,17^. Studies have reported the manipulation of this virus for oncolytic purposes ^18^. Use of engineered viruses is an emerging therapy rapidly evolving and holds a great promise as a potential treatment for various cancers. The current study aimed to elucidate the underlying mechanism of induced anti-proliferative effects by the engineered rMV-BNiP3 in breast tumor cells, particularly focusing on its selectivity for TNBC cells. The wild-type measles virus uses the SLAM receptor for entry into the host cells, while the Edmonston vaccine strain uses the CD46 receptor, which is generally overexpressed in various cancers, including breast cancer. It was found that the expression of CD46 was relatively higher in TNBC compared to non-TNBC cells. The modestly elevated expression of the CD46 receptor on TNBC cells may enhance the internalization of virus particles, potentially leading to improved anti-tumor effects. However, the difference is insignificant and it does not appear to be contributing to TNBC bias in treatment responses. Further research is needed to explore the interplay of other cellular pathways that influence the overall effectiveness of rMV-BNiP3 infection in TNBC cells. Recent studies had shown CD46 as a key player in both the transformation and treatment of cancers ^19^. The comprehensive analysis of the CD46 receptor and its role combined with the genetically modified rMV-BNiP3 targeting breast tumor cells is another aspect to be investigated.

Upregulation of β-catenin is commonly associated with various invasive tumors ^20^. It acts as a critical oncogenic driver in invasive breast cancer as well, particularly in TNBC/basal-like subtypes. Its nuclear accumulation of β-catenin promotes tumor cell proliferation, migration, invasion, and epithelial-mesenchymal transition (EMT) ^21^. Of the genes identified in our study on the basis of *in silico* analysis, the CTNNB1 gene encoding β-catenin of the Wnt/β-catenin pathway was identified as the main regulatory gene and the potential therapeutic target in breast tumor development. Wnt/β-catenin is a vital signaling pathway that plays multiple roles during embryonic development, cell migration, proliferation, differentiation, and survival ^22^. This signaling pathway is generally regulated well in all the developmental stages of the breast. It is believed that the activation of this aberrant pathway can enhance stem cell characteristics in TNBC cells ^23^. Reported studies and the extensive research suggest that the deregulation of this pathway could be a potential solution to minimize TNBC-related metastasis events associated with decreased stem cell properties ^24^. Moreover, this pathway is responsible for vasculogenic mimicry in TNBC cells supporting the importance of the Wnt/β-catenin signaling as a potential therapeutic target for future TNBC disease management.

Our experimental approaches, both *in silico* and *in vitro* studies, indicated that the rMV-BNiP3 targets β-catenin pathway subsequently the migratory ability of invasive TNBC cells. The downstream effectors including EGFR and Cyclin D1, are modulated by the c-JUN gene that encodes an oncoprotein (JUN) of critical importance in regulating genes promoting cell proliferation. c-JUN is often found to be overexpressed in many cancer subtypes, including nasopharyngeal carcinoma, mammary epithelial tumors, solid squamous cell carcinoma, and glioblastoma ^25^. Furthermore, another downstream target of β-catenin, matrix metalloprotease 7 (MMP-7) is known to play a role in cancer development and progression by regulating cancer cell proliferation, invasion, differentiation, and metastasis ^26^ in many cancer subtypes. These functions make cyclin D1 and MMP-7, potential targets for therapeutic interventions aimed at inhibiting tumor proliferation and migration.

In addition, the Wnt/β-catenin signal pathway is an important target to improve the efficacy of various cancer treatments. The activation of the Wnt/β-catenin pathway negatively affects the efficacy of chemotherapy. Long-term use of chemotherapy is associated with the development of resistance, which is linked to the increased activation of the Wnt/β-catenin signaling pathway ^27^. It was also reported that, β-catenin signaling in melanoma leads to resistance to immunotherapy, anti-PD-L1, and CTLA-4 monoclonal antibody therapy ^28^. Thus, the inhibition or reduction of β-catenin expression may sensitize the patients to chemotherapy and immunotherapy. As the engineered measles virus had demonstrated an inhibitory effect on β-catenin signaling, it may be administered in combination with chemotherapy, immunotherapy, and other anticancer drugs to enhance their therapeutic potential. Earlier, we also reported that the combination of chemotherapeutic agent, paclitaxel and rMV-BNiP3 resulted in significant TNBC cell death ^29^. It is evident from the current study that the invasive property of TNBC was greatly affected by the reduction of the β-catenin pathway and other proliferative pathways following rMB-BNiP3 infection.

In the current study and in our earlier study, anti-proliferative and migratory effects were observed to be promising in TNBC cells infected with rMV-BNiP3 compared to rMV lacking BNiP3, a proapoptotic gene in its genome. Though the contribution of BNiP3 to these effects is not clear, but it is hypothesised, that the engineered measles virus infection results in exogenously added BNiP3 expression, which may interact with Bcl2 as it has a Bcl2-interacting domain. The interaction between BNiP3 and Bcl2 allows Bax to release from the Bcl2-Bax complex, enabling Bax to induce apoptosis and other observed events. This hypothesis was supported by our preliminary in silico docking studies where the interaction between BNiP3 and Bcl2 was observed to be stronger and more stable than that of the Bax (Unpublished data).

Overall, this study provides valuable insights into therapeutic strategy of rMV-BNiP3 for treating metastatic breast cancers as standalone therapy or in combination with other conventional and targeted therapies. Further studies are ongoing to investigate the therapeutic potential of rMV-BNiP3 in preclinical settings using organotypic spheroids and xenograft nude mouse models. Additionally, evaluating the interaction of this therapy with existing treatment options will be crucial in developing an effective protocol for clinical use.

## Acknowledgements

Authors acknowledge Prof. Naidu Subbarao for his help in bioinformatics studies.

## Authors contributions: CRediT

MSR conceptualized, designed the experiments, supervised, data analysed, prepared the manuscript, funding acquired. AK carried out the TNBC/non-TNBC data curation, bioinformatics studies, analysis and experiments. KD and GSU carried out some experiments.

## Ethical approval

Not applicable

---

**Suppl Fig 1.**
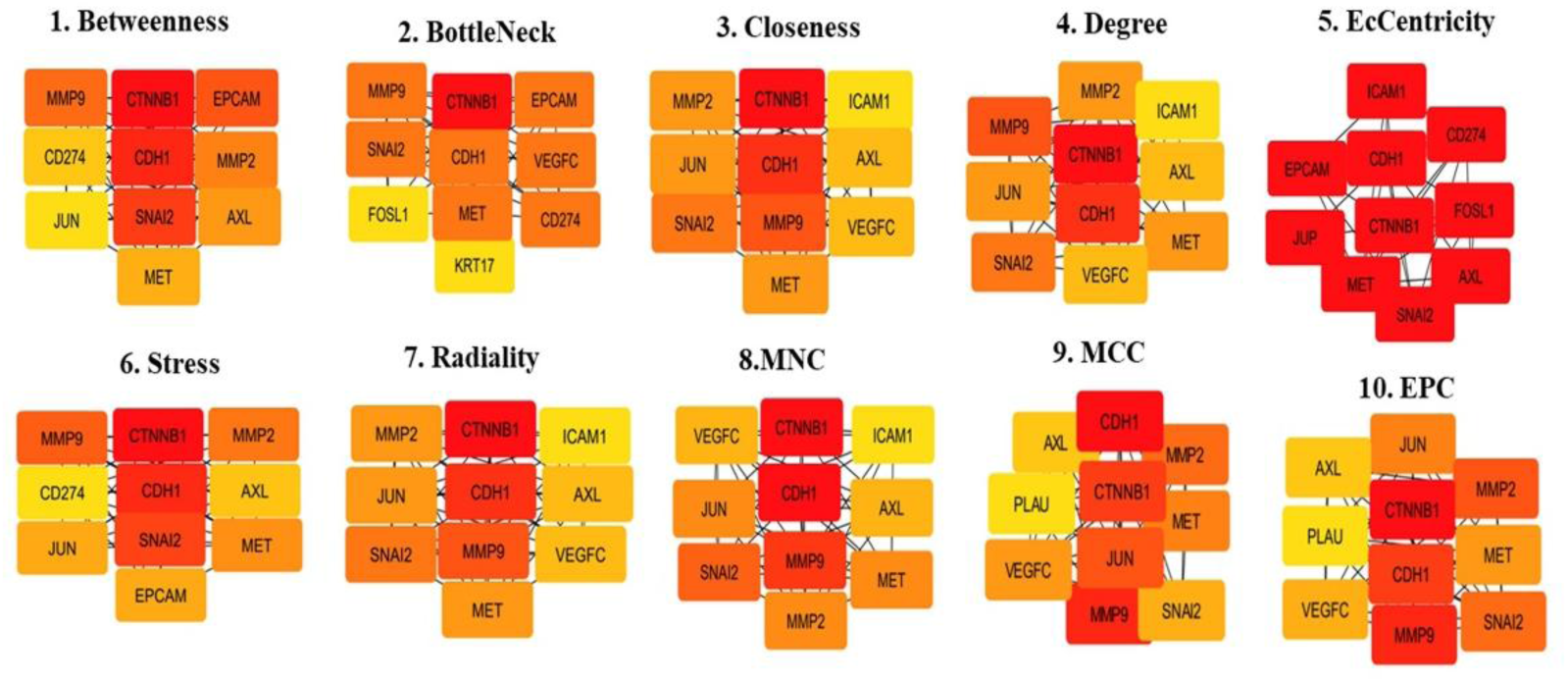
**Identification of hub genes**, key regulators of TNBC development and progression through PPI network using ten topological properties. Differentially expressed hub genes between MDA-MB-231 (representative TNBC cell line) *vs*. MCF-7 (representative of non-TNBC cell line).

